# MetaGxData: Clinically Annotated Breast, Ovarian and Pancreatic Cancer Datasets and their Use in Generating a Multi-Cancer Gene Signature

**DOI:** 10.1101/052910

**Authors:** Deena M.A. Gendoo, Michael Zon, Vandana Sandhu, Venkata SK Manem, Natchar Ratanasirigulchai, Gregory M Chen, Levi Waldron, Benjamin Haibe-Kains

## Abstract

A wealth of transcriptomic and clinical data on solid tumours are under-utilized due to unharmonized data storage and format. We have developed the *MetaGxData* package compendium, which includes manually-curated and standardized clinical, pathological, survival, and treatment metadata across breast, ovarian, and pancreatic cancer data. *MetaGxData* is the largest compendium of curated transcriptomic data for these cancer types to date, spanning 86 datasets and encompassing 15,249 samples. Open access to standardized metadata across cancer types promotes use of their transcriptomic and clinical data in a variety of cross-tumour analyses, including identification of common biomarkers, establishing common patterns of co-expression networks, and assessing the validity of prognostic signatures. Here, we demonstrate that *MetaGxData* is a flexible framework that facilitates meta-analyses by using it to identify common prognostic genes in ovarian and breast cancer. Furthermore, we use the data compendium to create the first gene signature that is prognostic in a meta-analysis across 3 cancers. These findings demonstrate the potential of *MetaGxData* to serve as an important resource in oncology research and provide a foundation for future development of cancer-specific compendia.

## Introduction

Ovarian, breast and pancreatic cancers are among the leading causes of cancer deaths among women, and recent studies have identified biological and molecular commonalities between them^1–4^. These cancers are part of hereditary syndromes related to mutations in a number of shared susceptibility genes that contribute to their carcinogenesis, including *BRCA1* and *BRCA2^3,5^*. As evidenced by epidemiological and linkage analysis studies, mutations and allelic loss in the *BRCA1* locus confers susceptibility to ovarian, pancreatic and early-onset breast cancer.^5–8^ The *BRCA2* gene appears to account for a proportion of early-onset breast cancer that is roughly equal to that resulting from *BRCA1*.^5,8^ *BRCA2*-mutation carriers with mutations within the ovarian cancer cluster region have been observed to exhibit greater risk for ovarian cancer.^5^ In addition to common susceptibility genes, both tumours may express a variety of common biomarkers that include hormone receptors, epithelial markers (e.g., cytokeratin 7, Ber-EP4), growth factor receptors (Her2/neu) and other surface molecules. ^3^

Commonalities between breast, ovarian, and pancreatic cancers have been observed not only for specific susceptibility genes, but at system-wide levels as well. In particular, molecular profiling across transcriptomes, copy-number landscapes, and mutational patterns emphasize strong molecular commonalities between basal-like breast tumours, high-grade serous ovarian cancer (HG-SOC), and basal-like pancreatic adenocarcinomas (PDACs).^2,9,10^ The growing list of parallels between Basal-like breast cancer, HG-SOC and basal-like PDACs include high frequency of *TP53* mutations and *TP53* loss, chromosomal instability, and widespread DNA copy number changes ^2,9–11^. Statistically significant subsets of both Basal-like breast tumors and HG-SOC also share *BRCA1* inactivation, *MYC* amplification, and highly correlated mRNA expression profiles.^2,9^ Subtype-specific prognostic signatures also reveal strong similarities between prognostic pathways in basal-like cancer and ovarian cancer, while ER-negative and ER-positive breast cancer subtypes exhibit different prognostic signatures ^12^. These ongoing studies promote identification of shared prognostic and predictive biomarkers across multiple cancer subtypes for future treatment.

Continuous growth of publicly available databases of breast, ovarian and pancreas genome-wide profiles necessitates the development of large-scale computational frameworks that can store these complex data types, as well as integrate them for meta-analytical studies. Current bioinformatics initiatives provide extensive data repositories for microarray data retrieval and annotation of specific tumour types. These resources enable analysis of single datasets, but do not provide sufficient standardization across independent studies of single or multiple cancer types ^13–19^ allow meta-analysis or other holistic analyses. This poses a challenge for meta-analytical investigations that aim to address global patterns across multiple forms of cancer, including for example, building multi-cancer gene signatures that generalize to new data^9,20,21^. Identifying robust prognostic signatures from transcriptomic data remains a major obstacle ^9,12,21^, and requires large sample sizes that can only be provided by large-scale meta-analysis ^20,22–26^. Additionally, most gene signatures derived from a single or small set of datasets are not generalizable to new data. In our recent systematic validation of ovarian signatures, primarily built from single datasets, we demonstrated that the concordance index of the best ovarian signatures only ranged from 0.54 to 0.58 ^27^, whereas a signatures trained by meta-analysis could improve significantly on this performance ^28^. The resulting standardized database of ovarian cancer profiles ^29^ enabled numerous subsequent meta-analyses and the development of statistical methodology. Efforts to standardize analyses of the transcriptomes of multiple cancer types have focused on coupling microarray repositories with graphical user interfaces to allow researchers to address targeted biologic questions on collective transcriptome datasets ^30–32^; however, these tools lack the generality to apply novel and potentially complex analyses.

An integrative framework is thus needed to harness the breadth of transcriptomic and clinical data from multiple cancer types, and to serve as a resource for integrative analysis across these aggressive cancer types. There are growing efforts towards the development of curated and clinically relevant microarray repositories for breast cancer, ovarian cancer, and pancreatic cancer data ^4,29,33–36^. These studies provide a solid foundation for the development of a controlled language for clinical annotations and standardized transcriptomic data representation across the three cancer types. Here, we have developed the *MetaGxData* package compendium, which includes manually-curated and standardized clinical, pathological, survival, and treatment metadata for breast, ovarian, and pancreatic cancer transcriptome data. *MetaGxData* is the largest, standardized compendium of breast, ovarian and pancreas microarray data to date, spanning 86 datasets and encompassing 15,249 samples. Standardization of metadata across these cancer types promotes the use of their expression and clinical data in a variety of cross-tumour analyses, including identification of common biomarkers, establishing common patterns of co-expression networks, assessing the validity of prognostic signatures, and the identification of new consensus signatures that reflects upon common biological mechanisms. In this paper, we present our flexible framework, unified nomenclature, as well as applications that demonstrate the analytical power of integrative analysis of a large number of breast, ovarian, and pancreatic cancer transcriptome datasets. As an example of its application, we integrated breast and ovarian cancer data to develop a multi-cancer gene signature and assessed its prognostic value in pancreatic cancer, demonstrating the existence of a multi-cancer prognostic gene signature.

## Results

### MetaGxData characterization and curation

The *MetaGxData* compendium integrates three packages containing curated and processed expression datasets for breast (*MetaGxBreast*), ovarian (*MetaGxOvarian*), and pancreatic (*MetaGxPancreas*) cancers. Our current framework extends upon the standardized framework we had already generated for curatedOvarianData ^29^. Our proposed enhancements facilitate rapid and consistent maintenance of our data packages as newer datasets are added, and provides enhanced user-versatility in terms of data rendering across single or multiple datasets (Fig. 1). All of these datasets can be downloaded through the MetaGxBreast, MetaGxOvarian and MetaGxPancreas R data packages publicly available through the Bioconductor ExperimentHub ^37^. Vignettes outlining how to access the MetaGxBreast, MetaGxOvarian and MetaGxPancreas datasets in R are available through the Bioconductor website.

**Figure 1:**
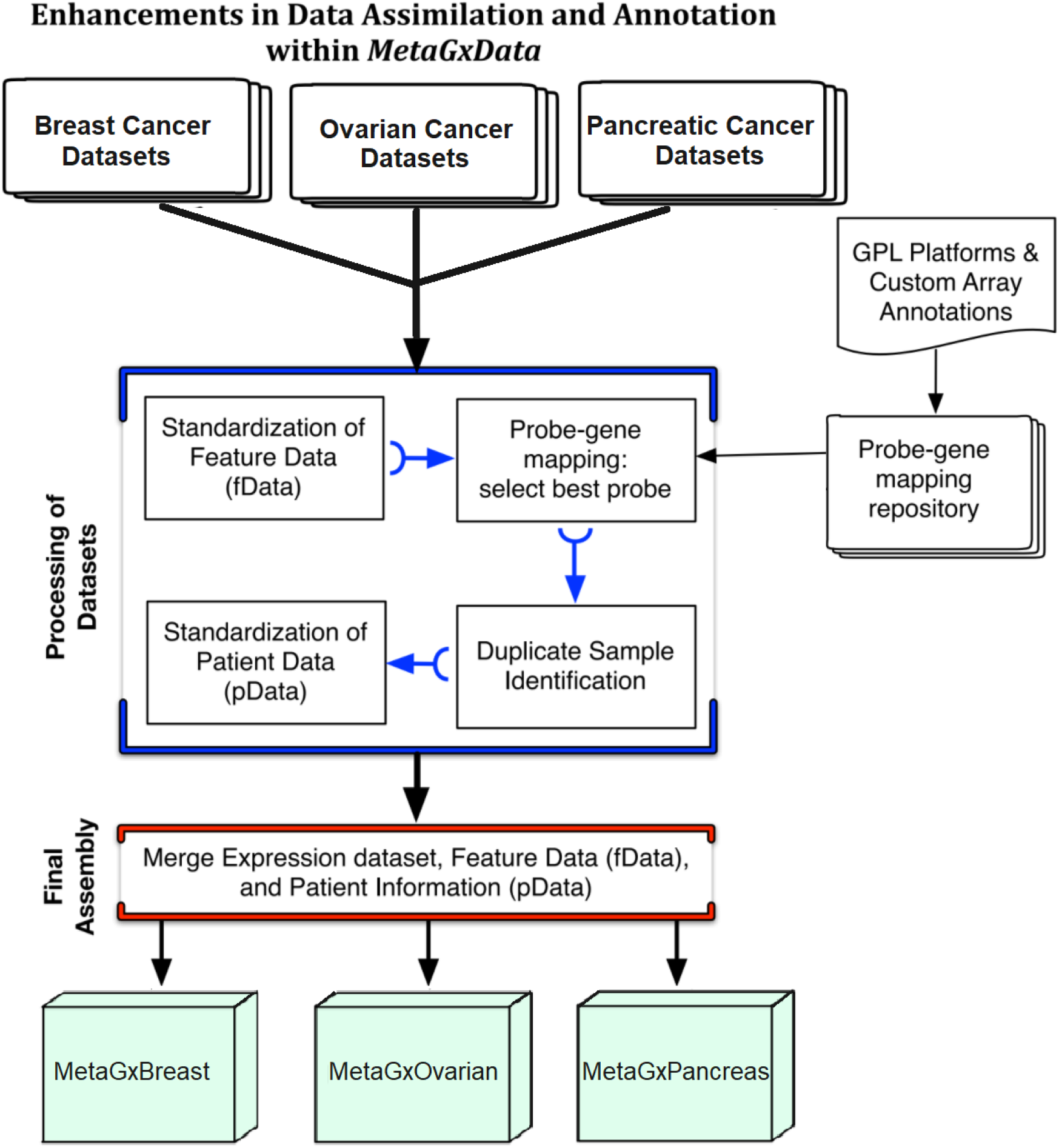
Diagrammatic representation of the enhancements in data integration and annotation within the MetaGxData framework. The process of downloading a dataset, and subsequent curation, annotation and integration into MetaGxData is depicted.

We developed semi-automatic curation scripts to standardize gene and clinical annotations of our breast, ovarian and pancreatic cancer datasets based on the nomenclature used in TCGA (Supplementary File S1) ^2,29^. Such annotations include a host of relevant categorical variables that reflect upon tumour histology (stage, grade, primary site, etc.), as well as categorical and numerical variables crucial for survival analysis and prognostication in these cancers (overall survival, recurrence-free survival, distant-free survival, and metastasis-free survival) (Supplementary Fig. S2). Most importantly, we have provided a number of comparable and overlapping clinicopathological features across breast, ovarian and pancreatic cancer samples, such as age at diagnosis, tumour grade, or vital status (Fig. 2). Additional common variables between the datasets can be seen in the supplementary figures (Supplementary Fig. S3, S4, S5). We also provide tumour-specific and critical annotations for each tumour type, including, for example, biomarker identification status (HER2, ER, PR) in breast cancer, and TNM status for pancreatic datasets. Treatment information across the cancers is provided when available, and survival information is focused exclusively on overall survival for pancreatic cancer.

**Figure 2:**
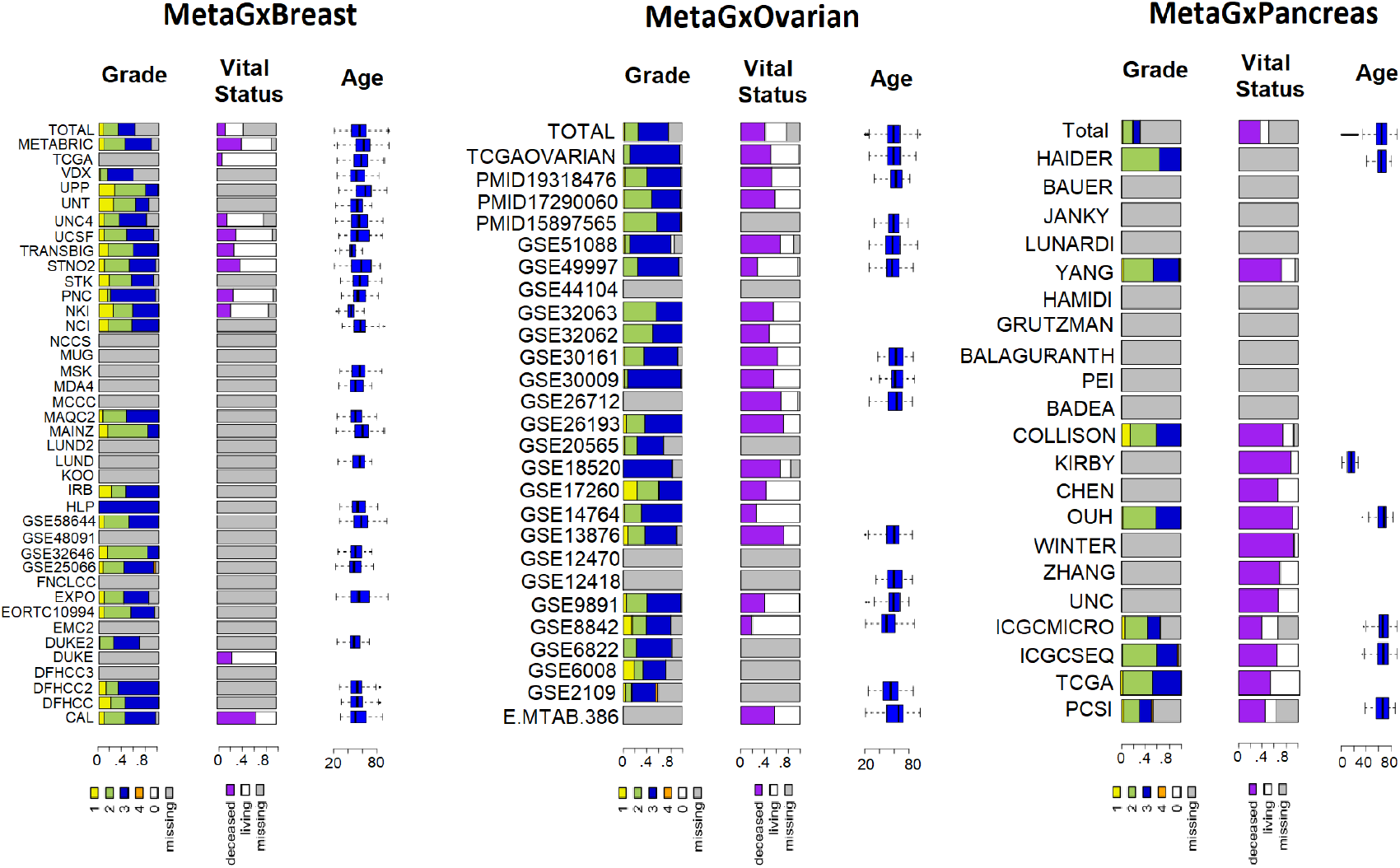
Schematic representation of some of the common clinical variables (pData) that are available across gene expression datasets in MetaGxBreast, MetaGxOvarian, and MetaGxPancreas. The Stacked bar plots indicate the percentage of samples in every dataset annotated with a particular variable designation. Continuous numeric values are represented by box plots.

### Analysis of prognostic genes in breast and ovarian cancer

The wealth and breadth of transcriptomic datasets in *MetaGxData* can be used as a framework for translational cancer research. As an example of the versatility of our packages, we conducted a meta-analysis of the prognostic value of well-studied prognostic genes in ovarian cancer and our previously published gene modules in breast cancer using the *MetaGxBreast* and *MetaGxOvarian* packages (Fig. 3, Fig. 4). The hazard ratio of the genes was determined by calculating the D.index, which is an estimate of the log hazard ratio comparing two equal sized groups. Furthermore, log rank tests were used to determine whether splits in the survival curves generated by using the genes to group patients into high and low score groups were statistically significant. A total of 13 genes and gene modules were tested, including 7 breast cancer gene modules (ESR1, ERBB2, STAT1, CASP3, PLAU, VEGF, and AURKA) and 6 ovarian cancer genes (PTCH1, TGFBR2, CXCL14, POSTN, FAP, and NUAK1).^22,23,27,28^ Unsurprisingly, higher gene expression levels of the proliferation gene AURKA indicate poorer survival in breast cancer (log rank p = 1.1e-16, n = 4,161) (Fig. 3c). This supports previous findings regarding the importance of this gene in biology-driven signatures of breast cancer, and its comparable prognostic effect with other multi-gene prognostic signatures.^22,23,35,38,39^ We have also observed that the NUAK1 gene exhibits worst prognosis in ovarian cancer (log rank p = 6.2e-9, n = 2,450) (Fig. 4c). We have previously demonstrated the utility of NUAK1 in the development of a debulking signature that can predict the outcome of cytoreductive surgery. ^28^

**Figure 3:**
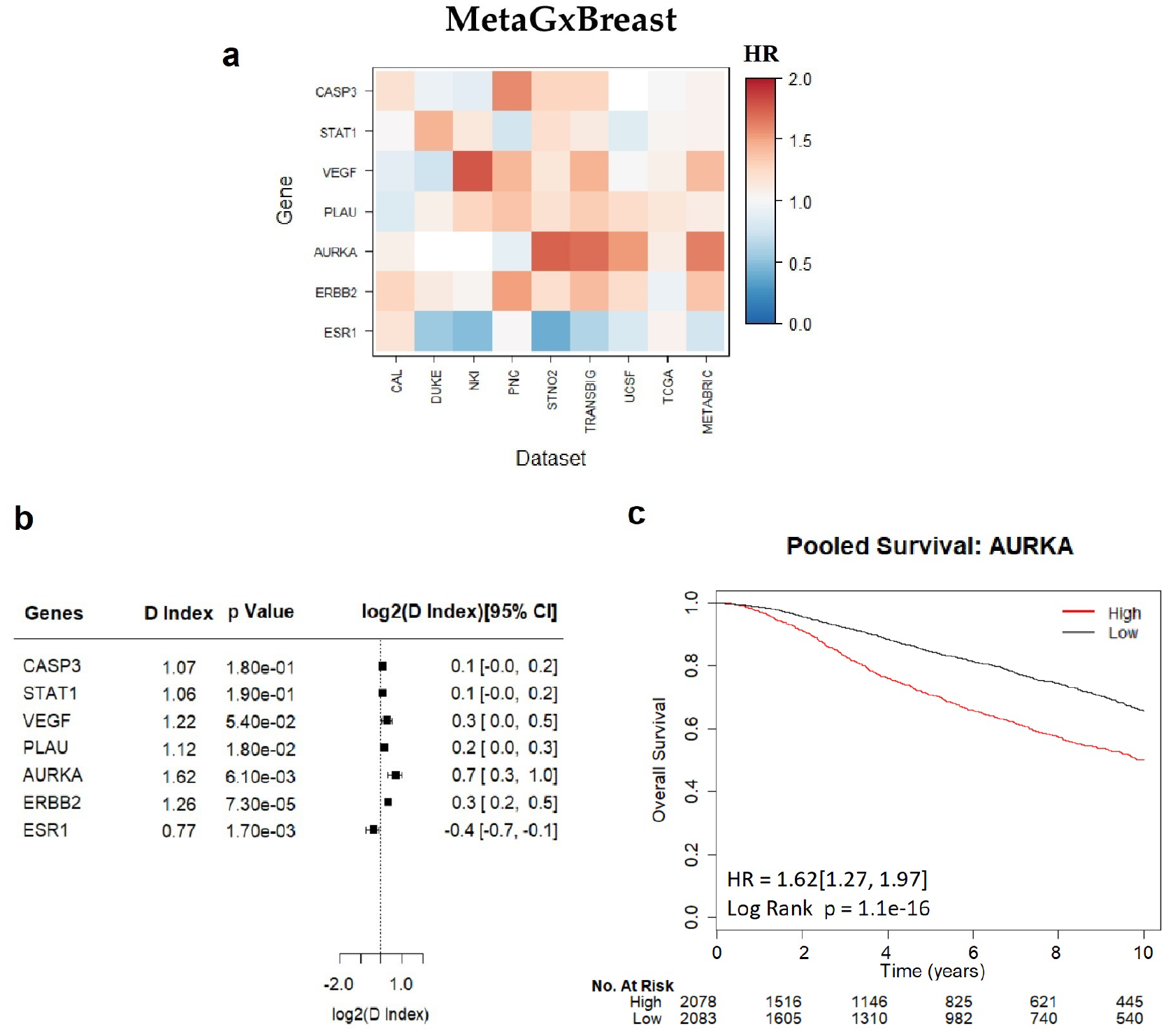
Assessment of the prognostic value of seven key gene modules in breast cancer, using the MetaGxBreast package. (A) Heatmap representation hazard ratios for each gene module, across 9 gene expression datasets. The estimate is presented as a hazard ratio for each gene. Ratios greater than 1 (red) indicate worse prognosis for elevated expression levels of that gene in the respective datasets. (B) Random effects meta-estimates of the hazard ratios for each gene, calculated by pooling the hazard ratios from each individual gene expression dataset. (C) Kaplan-Meier curve of the most prognostic gene with p < 0.05, in this case AURKA.

**Figure 4:**
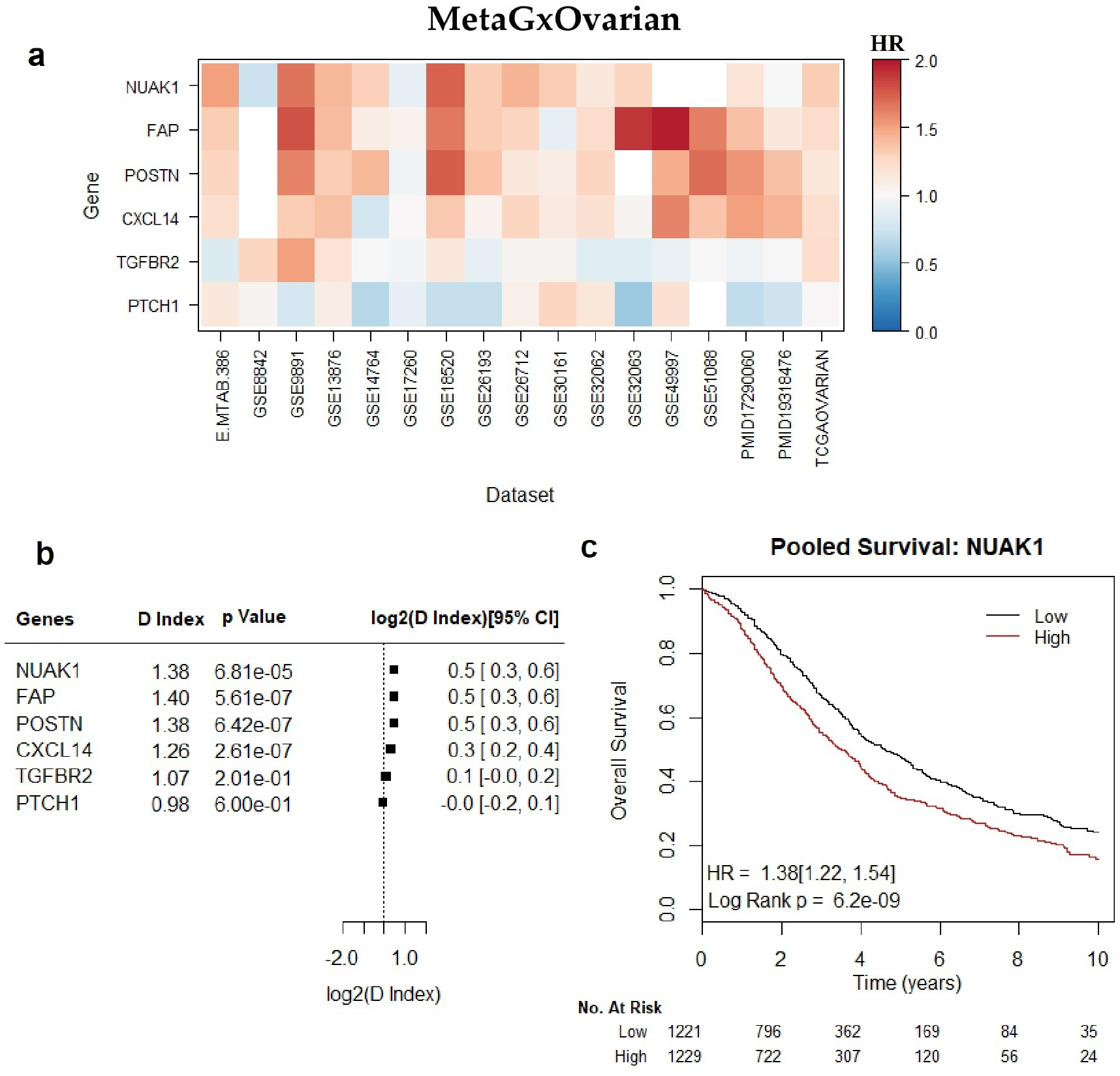
Assessment of the prognostic value of six key genes in ovarian cancer, using the MetaGxO-varian package. (A) Heatmap representation of hazard ratios for each gene, across 17 gene expression datasets. The estimate is presented as a hazard ratio for each gene. Ratios greater than 1 (red) indicate worse prognosis for elevated expression levels of that gene in the respective datasets. (B) Random effects meta-estimates of the hazard ratios for each gene, calculated by pooling the hazard ratios from each individual gene expression dataset. (C) Kaplan-Meier curve of NUAK1.

### Meta-analysis of gene expression prognosis across breast and ovarian cancer

Our single-gene prognostic analysis can easily be extended to a genome-wide meta-analysis. To this end, we determined the prognostic capability of 22,410 genes that are common to both the ovarian and breast cancer datasets (Fig. 5, Supplementary File S6). We identified 30 genes that are significantly prognostic across both tumours (False Discovery Rate [FDR] < 5%). Of these, we identified 3 genes for which elevated expression values indicate worse prognosis in both cancers (HR>1), and 9 genes for which it indicates better prognosis (HR<1)., Such analyses can be used to test pan-cancer hypotheses across much larger sample sizes than previously possible, and will allow deeper study of relationships between cancer subtypes such as will be integral in future studies of parallels between cancer subtypes, such as those comparing basal-like breast cancer, high-grade serous ovarian cancer (HG-SOC), and basal-like pancreatic cancer ^40^.

**Figure 5:**
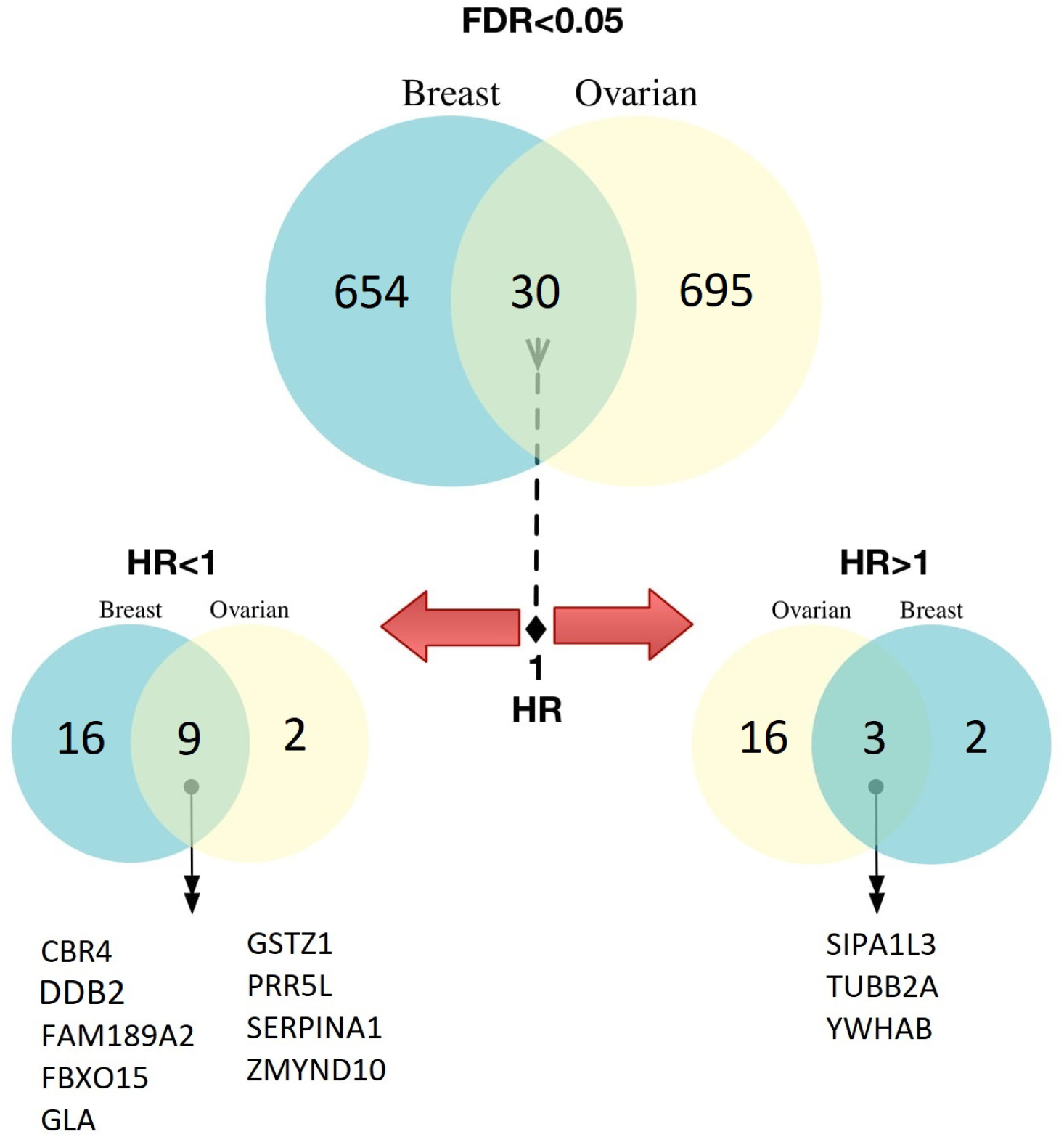
Genome-wide assessment of the prognostic value of 22,410 genes common to both the MetaGxBreast and MetaGxOvarian datasets. A Venn diagram of significant genes (FDR<5%) in each tumour following calculation of the Hazards Ratio is indicated (top). A total of 695 and 654 significantly prognostic genes were identified for ovarian and breast cancer, respectively. Common significant genes between both tumour types (n=30) were further subdivided by their log hazard ratio, for each tumour type. Genes for which elevated expression levels are prognostic (HR>1) across both tumours, or genes for which down-regulated expression is prognostic (HR<1) are indicated.

### MetaGx gene signature creation and prognosis in breast, ovarian and pancreatic cancer

We developed a gene signature that is prognostic in both breast and ovarian cancers by running a single-gene, genome-wide prognostic analysis on 22,410 genes as above, but excluding several large breast and ovarian datasets for use as validation cohorts. Meta-analysis of the training datasets identified 53 genes with significant hazard ratios in both cancers (FDR < 5%, HR > 1.125 or HR < 0.875) which were used to form the MetaGx signature (Table 1). The direction of association of the genes comprising the signature were chosen based on the hazard ratios (HR > 1 positive direction). The top 5 signatures from our recent review of ovarian gene signatures were evaluated alongside the MetaGx signature, and each signature was tested in the molecular subtypes identified by The Cancer Genome Atlas Research Network (immunoreactive, proliferative, mesenchymal, differentiated subtypes) ^1,27^. The MetaGx signature was the most prognostic of the ovarian signatures tested in an analysis containing all the patients (HR 2.02, n = 1,069) and was the only signature providing statistically significant prognostic capabilities within each subtype (log rank tests p < .05). Although the D index was prognostic in the differentiated subtype (HR 1.85, n = 427) and the most prognostic of the signatures tested in the Mesenchymal subtype (HR 1.95, n = 229), the MetaGx signature did not yield statistically significant D indices in the immunoreactive and proliferative subtypes (Fig. 6a-e).

**Figure 6:**
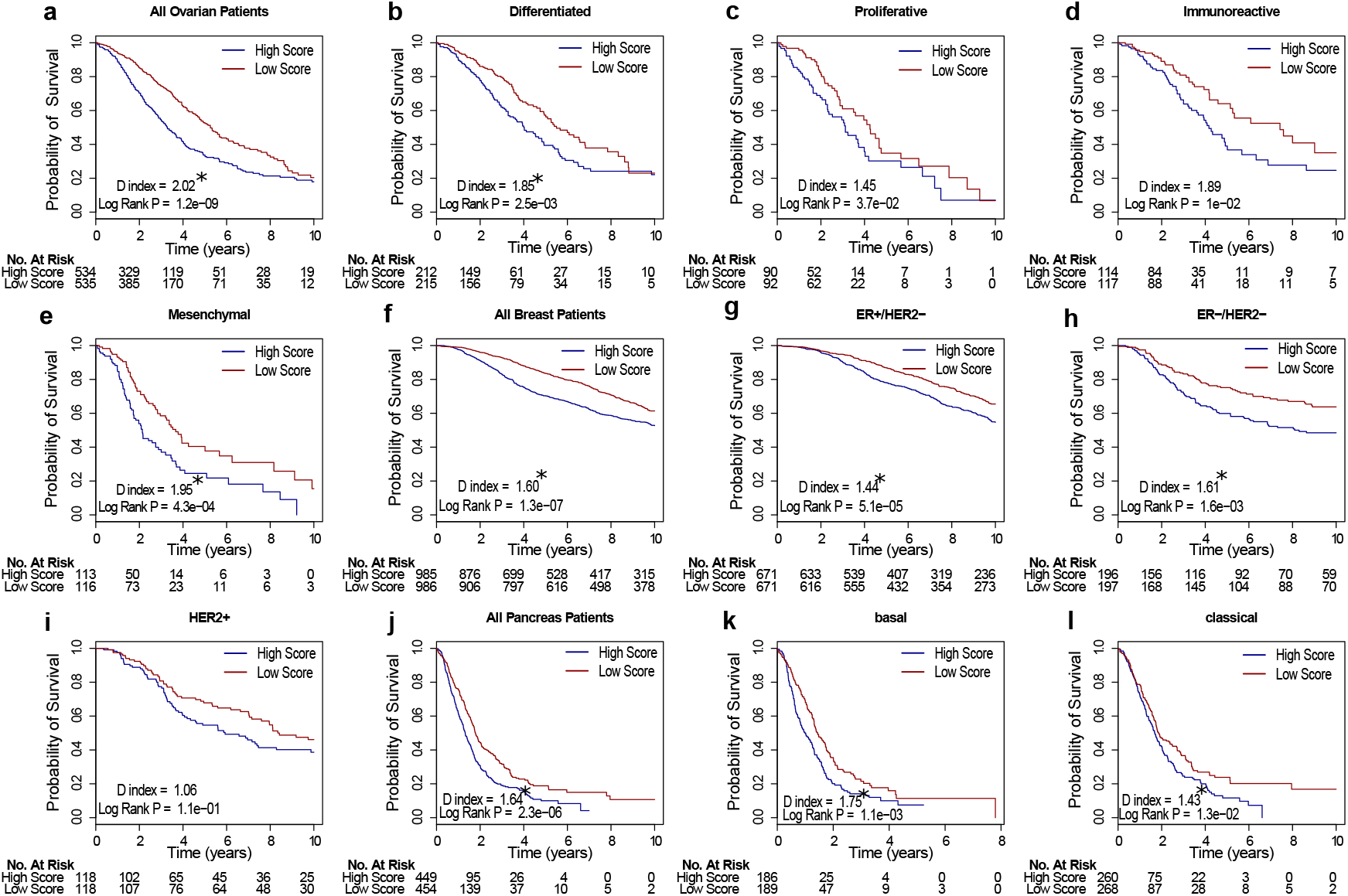
Survival curves for the MetaGx signature with patients stratified by molecular subtypes. (a-e) Survival curves in ovarian cancer. (f-i) Survival curves in breast cancer. (j-l) Survival curves in pancreatic cancer. The asterisks above the D indices indicate whether the D index was statistically significant (p < 0.05).

**Table 1:**
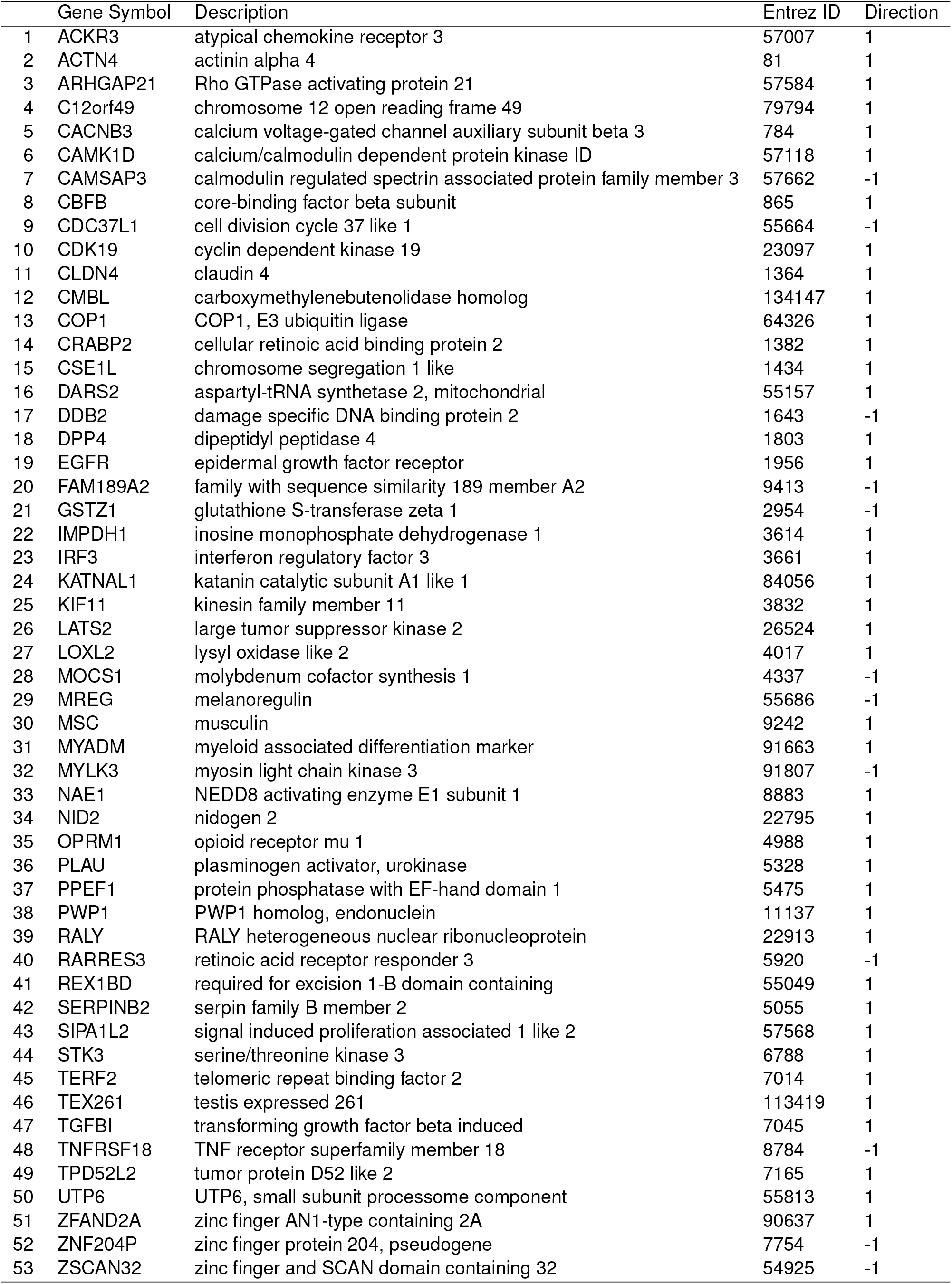
Genes present in the MetaGx gene signature.

In breast cancer, the signature was benchmarked against the clinically relevant mammaprint and oncotype DX signatures.^41–43^ Our three gene (ER, HER2, and AURKA) subtype classification model (SCM) was chosen to classify patients into the ER+/HER2-, ER-/HER2-, and HER2+ subtypes^35^. The MetaGx signature was highly prognostic in the analysis using all patients (HR 1.60, n = 1,971) (Fig. 6f) and had the largest D index in the ER-/HER2-subtype (HR 1.61, n = 393) (Supplementary Fig. S7).

We further tested the prognostic value of the MetaGx signature in pancreatic cancer and benchmarked it against pancreatic signatures from the literature, using a signed average approach for evaluation. ^44–47^ Of the 5 signatures tested, the MetaGx signature was the most prognostic in the analysis of all the patients (HR 1.64, n = 903) and was the only signature that yielded a statistically significant difference in survival within both the basal (log rank p = 1.1e-3, n = 375) and the classical (log rank p = 1.3e-2, n = 528) pancreatic cancer molecular subtypes identified by Moffitt et al (Table 2, Fig. 6j-l).^48^ Recent studies have shown that most published gene signatures often perform no better than 1000 random signatures of equal length. To test this observation, the MetaGx signature was tested in the pancreatic cancer, ovarian cancer and breast cancer test datasets against 1000 random signatures of equal size.^49^ In all three cases, the magnitude of the hazard ratio from the MetaGx signature was larger than the random signatures’ hazard ratio (p = 0.001 for all three cancers) (Supplementary Fig. S8).

**Table 2:** Prognostic value of pancreatic cancer gene signatures.

|  | Gene Signature - Subtype | D Index | D Index 95% CI | D Index P | Log Rank Test P |
| --- | --- | --- | --- | --- | --- |
| 1 | MetaGx - All Patients | 1.64 | (1.37, 1.90) | 1.9e-04 | 2.3e-06 |
| 2 | MetaGx - basal | 1.75 | (1.31, 2.19) | 1.1e-02 | 1.1e-03 |
| 3 | MetaGx - classical | 1.43 | (1.09, 1.77) | 3.7e-02 | 1.3e-02 |
| 4 | Newhook PLos one - All Patients | 1.22 | (0.97, 1.47) | 1.1e-01 | 1.2e-01 |
| 5 | Newhook PLos one - basal | 1.01 | (0.80, 1.23) | 9e-01 | 7.9e-01 |
| 6 | Newhook PLos one - classical | 0.99 | (0.77, 1.20) | 9e-01 | 6.2e-01 |
| 7 | Haider Gen Med - All Patients | 1.56 | (1.23, 1.88) | 6.8e-03 | 4.7e-06 |
| 8 | Haider Gen Med - basal | 1.22 | (0.88, 1.57) | 2.4e-01 | 2.2e-01 |
| 9 | Haider Gen Med - classical | 1.43 | (1.15, 1.71) | 1e-02 | 9.5e-02 |
| 10 | Grutzmann Oncogene - All Patients | 1.35 | (1.22, 1.49) | 1.3e-05 | 2.1e-06 |
| 11 | Grutzmann Oncogene - basal | 1.29 | (0.91, 1.67) | 1.7e-01 | 7.8e-03 |
| 12 | Grutzmann Oncogene - classical | 1.23 | (1.01, 1.46) | 6.2e-02 | 1.1e-01 |
| 13 | Stratford PLos med - All Patients | 1.39 | (1.09, 1.68) | 2.9e-02 | 6.3e-03 |
| 14 | Stratford PLos med - basal | 1.22 | (1.01, 1.43) | 6.4e-02 | 2.6e-01 |
| 15 | Stratford PLos med - classical | 1.29 | (0.94, 1.63) | 1.4e-01 | 6.7e-02 |

## Discussion

Meta-analysis of multiple cancer types is an area of high interest, with ongoing research continually supporting the growing relationship between these malignancies and suggesting common patterns of tumour biology ^50^. We provide an integrative, standardized, and comprehensive platform to facilitate analysis between breast, ovarian, and pancreatic cancer. This platform provides a flexible framework for data assimilation and unified nomenclature, with standardized data packages hosting the largest compendia of breast, ovarian, and pancreatic cancer transcriptomic and clinical datasets available to date.

Integration of genomic data into standardized frameworks is challenged by the inconsistency of the clinical curations across datasets and across tumour types. Annotation of clinicopathological variables may vary widely due to different protocols in different laboratories, institutions, and across international boundaries. We have standardized, as much as possible, the catalog of clinical variables within each tumour type. For characteristics pertaining to a specific tumour type, including ER, PGR, and HER2 IHC status in breast cancer samples, we have generated a semantic positive/negative variable to reflect IHC status. This facilitates searching across all patients irrespective of the original assay annotations that may have binary, numeric, or qualitative. Similarly, a binary variable has been assigned to ovarian cancer patients to reflect whether they had been treated with platinum, taxol, or neoadjuvant therapy. Many of the annotated variables (ex: stage and tumour grade in MetaGxOvarian) have also been standardized to facilitate comparisons across multiple studies. Further analyses using our previously developed packages (curatedOvarianData) have indicated good consistency across datasets, and ultimately facilitated uniform and consistent investigations on the prognostic effect of biomarkers in ovarian cancer survival ^51,52^.

The scale of MetaGxData facilitates identification of gene signatures that are prognostic across multiple forms of cancer. Using this compendium, we developed a gene signature that is prognostic for breast, ovarian, and pancreatic cancers. Requiring genes to be prognostic across multiple datasets should help distinguish between general and disease-specific processes affecting patient survival. allow signatures to generalize better to new datasets, in comparison to conventional signature creation methods that select genes based on cox proportional hazard models in a single dataset. We have demonstrated that the This multi-cancer MetaGx signature outperformed the top ovarian signatures identified in our previous review in an analysis conducted on all patients with overall survival as the endpoint. It was also more prognostic than the clinically-relevant Mammaprint and OncotypeDX signatures in the ER-/HER2-breast cancer subtype, and more prognostic than pancreas-specific signatures in pancreatic cancer. Furthermore, it was the only signature that was prognostic in each molecular subtype of pancreatic cancer, and was highly prognostic in the basal-like subtype

To our knowledge, the MetaGx signature represents the first signature demonstrated to be prognostic in a meta-analysis across three cancers. This includes pancreatic cancer, which was not used in any way for training. This signature includes notable genes such as PLAU, which we have previously shown to be associated with tumor invasion/metastasis, as well as epidermal growth factor receptor.^23,53^ Our signature provides additional support for the role of CLDN4 in pancreas, breast and ovarian malignancies. Higher expression levels of this gene placed patients in the high score group that had poorer outcomes in all 3 of these cancers. This is in line with numerous studies that have shown CLDN4 to be overexpressed in pancreatic, ovarian, and breast tumors relative to normal tissue.^54–58^

In conclusion, the MetaGxBreast, MetaGxOvarian and MetaGxPancreas packages follow a unified framework that facilitates integration of oncogenomic and clinicopathological data. We have demonstrated how our packages facilitate easy meta-analysis of gene expression and prognostication in breast, ovarian and pancreatic cancer. We have also demonstrated that leveraging this data in meta-analysis can lead to gene signatures that outperform clinically relevant breast signatures in ER-/HER2-patients, top ovarian signatures developed from single datasets, and a number of published pancreatic cancer signatures. These packages have the potential to serve as an important resource in oncology and methodological research and provide a foundation for future development of cancer-specific compendia.

## Methods

### Breast cancer data acquisition

Breast cancer datasets were extracted from our previous meta-analysis of breast cancer molecular subtypes, which includes 39 microarray datasets from a variety of commercially available microarray platforms published from 2002 to 2014 ^35^. Additional datasets were extracted from the Gene Expression Omnibus (GEO) and manually curated. Gene expression and clinical annotation for Metabric were downloaded from EBI ArrayExpress and combined into a dataset of 2,136 samples ^59^. The cgdsr R package was used to extract 1,098 tumour samples from The Cancer Genome Atlas (TCGA), and matching clinical annotations for these samples were downloaded from the TCGA Data Matrix portal (https://tcga-data.nci.nih.gov/tcga/) ^2,60^. Combining these studies produced a total of 39 breast cancer microarray expression datasets spanning 10,004 samples. Of these 10,004 samples, survival information is available for 6,847 patients, including overall survival (n=4425), metastasis free survival (n=2695), and relapse free survival (n=1858).

### Ovarian cancer data acquisition

Ovarian microarray expression datasets were obtained from our recent update of the curatedOvarianData data package, onto which we have added 5 expression datasets to the originally published version ^29^, for a total of 26 microarray datasets spanning 3,526 samples. To obtain these datasets we first used the curatedOvarianData pipeline to generate the “FULLcuratedOvarianData” version of the package, which differs from the public version in that probe sets for same gene are not merged (https://bitbucket.org/lwaldron/curatedovariandata). Of the 3,526 samples, survival information is available for 2,726 patients, including overall survival (n=2,,712) and relapse free survival (n=1928).

### Pancreatic cancer data acquisition

Pancreatic ductal adenocarcinoma (PDAC) datasets were obtained by curating datasets available from the literature. A total of 21 datasets were curated for a total of 1,719 patient transcriptomic profiles. Of the 21 datasets, overall survival data was present for 12 studies. Consequently, of the 1,719 samples survival information is available for 1000 patients, including overall survival (n=1000) and no relapse free survival data.

### Processing of gene expression datasets

The processing of breast and ovarian cancer microarray datasets was previously described ^29,35^. The pancreatic cancer datasets were processed in the manner described within the original studies from which they were obtained; the only exception is the Kirby dataset, which had been aligned using Kallisto and whose expression values are calculated using the logarithm of the transcripts per kilobase million (TPM). Across all datasets, we used GEO platform descriptions as the primary source of probe and gene annotations when available, otherwise original annotations as published by the authors were used for non-standard gene expression profiling platforms. The full set of gene annotation platforms across all expression sets can be found in the metadata files associated with each Bioconductor package. Gene symbols and Entrez Gene identifiers that matched the probeset ids of a given expression set were subsequently saved as part of the featureData (fData) pertaining to that expression set. For genes with multiple probesets, only the probe with the highest variance across the dataset was used to calculate the prognostic value of the gene.

### MetaGxData package implementation

The breast, ovarian, and pancreatic cancer datasets are available through the MetaGxBreast, MetaGxOvarian, and MetaGxPancreas R data packages hosted on Bioconductor’s ExperimentHub. The MetaGxData packages allow users to select and filter the finalized curated datasets using the loadOvarianEsets, loadBreastEsets and loadPancreasEsets functions of MetaGxOvarian, MetaGxBreast and MetaGxPancreas, respectively. Users are provided options for filtering samples based on clinical parameters, availability of survival data, and sample replicates (patients with highly correlated transcriptomic profiles; spearman correlation > 0.98). Users are also provided other options including, but not limited to, the ability to remove datasets based on the number of samples and the number of survival events present in the data. Importantly, users have the ability to specifically select for only primary tumour samples or several tissue types (primary tumours, healthy tissue, etc.) using the sample type info found in the clinical data.

Collectively, our data compendium, referred to as *MetaGxData*, encompasses 86 processed gene expression datasets, containing in total 15,249 breast, ovarian and pancreas samples. Information pertaining to the breast, ovarian, and pancreas datasets can be found in in the supplementary files (Supplementary Table S9, S10, and S11). Expression datasets are represented as SummarizedExperiment objects with attached clinical data (pData), and feature data (fData) and can be loaded into R with a single function call allowing for fast and flexible analysis.61 Hosting the datasets within the Bioconductor ExperimentHub facilitates rapid integration of new datasets into the existing framework and allows for easy extension of newer studies into the package in future iterations of *MetaGxData*.

### Prognostication of breast and ovarian cancer genes and signature generation

Cox proportional hazards analysis was performed using the R package *survcomp* (version 1.29.4) to estimate the prognostic value (hazard ratio) and significance (corresponding p-value) of the genes in the various datasets.^62^ Overall survival was used as the primary endpoint and meta-estimates of the hazard ratio were calculated using the random-effects model ^63^. When stratifying samples into groups to generate survival curves, samples within each dataset were stratified into two groups based on the median expression of the gene or the median gene signature/module score for all the samples within that dataset. For the gene signatures, risk prediction scores were determined using the signed average of the patients’ gene expression with the sign being determined as their direction of association with the survival outcome (HR > 1 positive direction). In order to generate the MetaGx gene signature, the aforementioned analysis was performed on the genes within MetaGxData while removing the METABRIC dataset (n=2136 samples) from MetaGxBreast, and 5 of the largest ovarian datasets (GSE9891, GSE32062, GSE49997, GSE26712, GSE51088, totalling 1,116 samples) for use as the validation cohort. The 53 genes with significant hazard ratios in both cancers (FDR < 5%, HR > 1.125 or HR < 0.875) were selected for the MetaGx gene signature

### Statistical analysis

Risk predictions for the signature along with the patients’ corresponding survival times and overall survival status were used in the R survcomp package in order to compute a D index in each dataset.^62,64^ Meta-analyses were also conducted via survcomp to obtain a single best estimate of the D index (random effects model) using the D indices computed for each individual dataset. The patient groups, survival times and overall survival status of the patients from all the datasets were used within the survival package in order to generate a Kaplan-Meir survival curve and determine the log-rank test p values.^65^ D index and log-rank test p values less than 0.05 were considered to be statistically significant and all analyses were conducted using R.

### Research reproducibility

All the code required to reproduce the single-gene prognosis analysis, as well as the genome-wide meta-analysis and signature results, is publicly available on the CodeOcean (codeocean.com, analysis at http://bit.ly/2Muk5io). The CodeOcean contains an executable version of the code, in the form of a standalone docker, that can be used to generate all of the results in the present work. This work complies with the guidelines outlined in ^66–68^.

## Data Availability

The datasets used in this manuscript are all publicly available for download through R Bioconductor’s ExperimentHub (https://bioconductor.org/packages/release/data/experiment/). The breast, ovarian, and pancreas datasets can be found in MetaGxbreast, MetaGxOvarian, and MetaGxPancreas, Respectively

## Supplementary Files

**Supplementary File S1.** Explanation of curated clinical annotations (phenotype data variables) in MetaGxBreast (sheet 1), MetaGxOvarian (sheet 2), and MetaGxPancreas (sheet 3).

**Supplementary Figure S2.** Heatmap representation of clinical variables availability across gene expression datasets of MetaGxBreast, MetaGxOvarian, and MetaGxPancreas. Datasets are represented as rows and clinical variables as columns. The percentage of samples in each dataset that is annotated with a particular variable is represented.

**Supplementary Figure S3.** Schematic representation of the clinical variables (pData) that are available across gene expression datasets in MetaGxBreast. The Stacked bar plots indicate the percentage of samples in every dataset annotated with a particular variable designation.

Continuous numeric values are represented by box plots.

**Supplementary Figure S4.** Schematic representation of the clinical variables (pData) that are available across gene expression datasets in MetaGxOvarian. The Stacked bar plots indicate the percentage of samples in every dataset annotated with a particular variable designation.

Continuous numeric values are represented by box plots.

**Supplementary Figure S5.** Schematic representation of the clinical variables (pData) that are available across gene expression datasets in MetaGxPancreas. The Stacked bar plots indicate the percentage of samples in every dataset annotated with a particular variable designation.

Continuous numeric values are represented by box plots.

**Supplementary File S6.** Genome-wide analysis of the prognostic value of 22,410 genes in breast and ovarian gene expression datasets. (sheet 1) List of the computed Hazard Ratio of all genes, using MetaGxBreast. (sheet 2) List of the computed Hazard Ratio of all genes, using MetaGxOvarian.

**Supplementary Figure S7.** Survival curves for the metaGx, mammaprint signature, and oncotype signature in the ER-/HER2-breast cancer patients stratified by molecular subtypes.

The asterisks above the D indices indicate whether the D index was statistically significant (p < 0.05).

**Supplementary Figure S8.** Density plots comparing the prognostic value of the gene signature in breast, ovarian, and pancreatic cancer to 1000 random signatures of the same length. The dashed line represents the location of the D index of the signature.

**Supplementary Table S9.** Breast cancer dataset information

**Supplementary Table S10.** Ovarian cancer dataset information

**Supplementary Table S11.** Pancreatic cancer dataset information

## Acknowledgements

The authors would like to thank all the authors who made available their valuable gene expression and clinical data for breast, ovarian, and pancreatic cancers over the past two decades. This study was conducted with the support of the Ontario Institute for Cancer Research (OICR, PanCuRx Translational Research Initiative) through funding provided by the Government of Ontario (Ministry of Research, Innovation, and Science). G.M. Chen was supported by a Computational Biology Undergraduate Summer Student Health Research Award. V.S was supported by grants from The Radium Hospital Foundation, Oslo University Hospital, and the PanCuRx Translational Research Initiative at the OICR. V.S.K.M was supported by the Cancer Research Society. B.H.K was supported by the Gattuso Slaight Personalized Cancer Medicine Fund at Princess Margaret Cancer Centre, the Canadian Institutes of Health Research, the Natural Sciences and Engineering Research Council of Canada, and the Ministry of Economic Development and Innovation/Ministry of Research & Innovation of Ontario (Canada). L.W. was supported by the National Cancer Institute at the National Institutes of Health (1R03CA191447-01A1).

## Author Contributions

D.M.A.G and N.R. designed and developed the processing pipeline for the MetaGxData framework, and developed the MetaGxBreast and MetaGxOvarian packages. V.S processed and curated the data present in MetaGxPancreas. M.Z organized the data and developed the MetaGxBreast, MetaGxPancreas, and MetaGxOvarian packages on Bioconductor. D.M.A.G, M.Z and N.R. conducted the single-gene prognosis and genome-wide single-gene analysis. M.Z. developed the gene signature and wrote the code that generates the manuscript’s results. M.Z. developed the CodeOcean capsule setting up a fully-specified docker container to reproduce all the analysis results. B.H.K designed and supervised the study.

## References

1. Cancer Genome Atlas Research Network. Integrated genomic analyses of ovarian carcinoma. Nature 474, 609–615 (2011).

2. The Cancer Genome Atlas Network. Comprehensive molecular portraits of human breast tumours. Nature 490, 61 (2012).

3. Davidson, B. et al. Gene expression signatures differentiate ovarian/peritoneal serous carcinoma from breast carcinoma in effusions. J. Cell. Mol. Med. 15, 535–544 (2011).

4. Chelala, C. et al. Pancreatic Expression database: a generic model for the organization, integration and mining of complex cancer datasets. BMC Genomics 8, 439 (2007).

5. Greer, J. B. & Whitcomb, D. C. Role of BRCA1 and BRCA2 mutations in pancreatic cancer. Gut 56, 601–605 (2007).

6. Futreal, P. A. et al. BRCA1 mutations in primary breast and ovarian carcinomas. Science 266, 120–122 (1994).

7. Billack, B. & Monteiro, A. N. A. BRCA1 in breast and ovarian cancer predisposition. Cancer Lett. 227, 1–7 (2005).

7. Ford, D. & Easton, D. F. The genetics of breast and ovarian cancer. Br. J. Cancer 72, 805–812 (1995).

9. Michiels, S., Koscielny, S. & Hill, C. Prediction of cancer outcome with microarrays: a multiple random validation strategy. Lancet 365, 488–492 (2005).

10. Sandhu, V. et al. The Genomic Landscape of Pancreatic and Periampullary Adenocarcinoma. Cancer Res. 76, 5092–5102 (2016).

11. Bailey, P. et al. Genomic analyses identify molecular subtypes of pancreatic cancer. Nature 531, 47–52 (2016).

12. Macgregor, P. F. Gene expression in cancer: the application of microarrays. Expert Rev. Mol. Diagn. 3, 185–200 (2003).

13. Cheng, W.-C. et al. Microarray meta-analysis database (M2DB): a uniformly pre-processed, quality controlled, and manually curated human clinical microarray database. BMC Bioinformatics 11, 421 (2010).

14. Coletta, A. et al. InSilico DB genomic datasets hub: an efficient starting point for analyzing genome-wide studies in GenePattern, Integrative Genomics Viewer, and R/Bioconductor. Genome Biol. 13, R104 (2012).

15. Edgar, R., Domrachev, M. & Lash, A. E. Gene Expression Omnibus: NCBI gene expression and hybridization array data repository. Nucleic Acids Res. 30, 207–210 (2002).

16. Kolesnikov, N. et al. ArrayExpress update--simplifying data submissions. Nucleic Acids Res. 43, D1113–6 (2015).

17. Reich, M. et al. GenePattern 2.0. Nat. Genet. 38, 500–501 (2006).

18. Wan, Q. et al. BioXpress: an integrated RNA-seq-derived gene expression database for pan-cancer analysis. Database 2015, (2015).

19. Kannan, L. et al. Public data and open source tools for multi-assay genomic investigation of disease. Brief. Bioinform. 17, 603–615 (2016).

20. Ein-Dor, L., Zuk, O. & Domany, E. Thousands of samples are needed to generate a robust gene list for predicting outcome in cancer. Proc. Natl. Acad. Sci. U. S. A. 103, 5923–5928 (2006).

21. Ein-Dor, L., Kela, I., Getz, G., Givol, D. & Domany, E. Outcome signature genes in breast cancer: is there a unique set? Bioinformatics 21, 171–178 (2005).

22. Wirapati, P. et al. Meta-analysis of gene expression profiles in breast cancer: toward a unified understanding of breast cancer subtyping and prognosis signatures. Breast Cancer Res. 10, R65 (2008).

23. Desmedt, C. et al. Biological processes associated with breast cancer clinical outcome depend on the molecular subtypes. Clin. Cancer Res. 14, 5158–5165 (2008).

24. Chen, G. M. et al. Consensus on Molecular Subtypes of High-grade Serous Ovarian Carcinoma. Clin. Cancer Res. clincanres.0784.2018 (2018).

25. doi:10.1101/355602

26. Fishel, I., Kaufman, A. & Ruppin, E. Meta-analysis of gene expression data: a predictor-based approach. Bioinformatics 23, 1599–1606 (2007).

27. Waldron, L. et al. Comparative meta-analysis of prognostic gene signatures for late-stage ovarian cancer. J. Natl. Cancer Inst. 106, (2014).

28. Riester, M. et al. Risk prediction for late-stage ovarian cancer by meta-analysis of 1525 patient samples. J. Natl. Cancer Inst. 106, (2014).

29. Ganzfried, B. F. et al. curatedOvarianData: clinically annotated data for the ovarian cancer transcriptome. Database 2013, bat013 (2013).

30. Wettenhall, J. M., Simpson, K. M., Satterley, K. & Smyth, G. K. affylmGUI: a graphical user interface for linear modeling of single channel microarray data. Bioinformatics 22, 897–899 (2006).

31. Kapushesky, M. et al. Expression Profiler: next generation–an online platform for analysis of microarray data. Nucleic Acids Res. 32, W465–W470 (2004).

32. Parkinson, H. et al. ArrayExpress–a public database of microarray experiments and gene expression profiles. Nucleic Acids Res. 35, D747–D750 (2007).

33. Madden, S. F. et al. BreastMark: an integrated approach to mining publicly available transcriptomic datasets relating to breast cancer outcome. Breast Cancer Res. 15, R52 (2013).

34. Planey, C. R. & Butte, A. J. Database integration of 4923 publicly-available samples of breast cancer molecular and clinical data. AMIA Jt Summits Transl Sci Proc 2013, 138–142 (2013).

35. Haibe-Kains, B. et al. A three-gene model to robustly identify breast cancer molecular subtypes. J. Natl. Cancer Inst. 104, 311–325 (2012).

36. Madden, S. F. et al. OvMark: a user-friendly system for the identification of prognostic biomarkers in publically available ovarian cancer gene expression datasets. Mol. Cancer 13, 241 (2014).

37. Pasolli, E. et al. Accessible, curated metagenomic data through ExperimentHub. Nat. Methods 14, 1023–1024 (2017).

38. Gendoo, D. M. A. et al. Genefu: an R/Bioconductor package for computation of gene expression-based signatures in breast cancer. Bioinformatics 32, 1097–1099 (2016).

39. Sotiriou, C. et al. Gene expression profiling in breast cancer: understanding the molecular basis of histologic grade to improve prognosis. J. Natl. Cancer Inst. 98, 262–272 (2006).

40. Bowtell, D. D. L. The genesis and evolution of high-grade serous ovarian cancer. Nat. Rev. Cancer 10, 803–808 (2010).

41. Cardoso, F. et al. 70-Gene Signature as an Aid to Treatment Decisions in Early-Stage Breast Cancer. N. Engl. J. Med. 375, 717–729 (2016).

42. Kuijer, A. et al. Impact of 70-Gene Signature Use on Adjuvant Chemotherapy Decisions in Patients With Estrogen Receptor-Positive Early Breast Cancer: Results of a Prospective Cohort Study. J. Clin. Oncol. 35, 2814–2819 (2017).

43. McVeigh, T. P. & Kerin, M. J. Clinical use of the Oncotype DX genomic test to guide treatment decisions for patients with invasive breast cancer. Breast Cancer 9, 393–400 (2017).

44. Newhook, T. E. et al. A thirteen-gene expression signature predicts survival of patients with pancreatic cancer and identifies new genes of interest. PLoS One 9, e105631 (2014).

45. Haider, S. et al. A multi-gene signature predicts outcome in patients with pancreatic ductal adenocarcinoma. Genome Med. 6, 105 (2014).

46. Grützmann, R. et al. Meta-analysis of microarray data on pancreatic cancer defines a set of commonly dysregulated genes. Oncogene 24, 5079–5088 (2005).

47. Stratford, J. K. et al. A six-gene signature predicts survival of patients with localized pancreatic ductal adenocarcinoma. PLoS Med. 7, e1000307 (2010).

48. Moffitt, R. A. et al. Virtual microdissection identifies distinct tumor- and stroma-specific subtypes of pancreatic ductal adenocarcinoma. Nat. Genet. 47, 1168–1178 (2015).

49. Venet, D., Dumont, J. E. & Detours, V. Most random gene expression signatures are significantly associated with breast cancer outcome. PLoS Comput. Biol. 7, e1002240 (2011).

50. Hanahan, D. & Weinberg, R. A. Hallmarks of cancer: the next generation. Cell 144, 646–674 (2011).

51. Cheng, X., Lu, W. & Liu, M. Identification of homogeneous and heterogeneous variables in pooled cohort studies. Biometrics 71, 397–403 (2015).

52. Trippa, L., Waldron, L., Huttenhower, C. & Parmigiani, G. Bayesian nonparametric cross-study validation of prediction methods. Ann. Appl. Stat. 9, 402–428 (2015).

53. Nicholson, R. I., Gee, J. M. & Harper, M. E. EGFR and cancer prognosis. Eur. J. Cancer 37 Suppl 4, S9–15 (2001).

54. Hewitt, K. J., Agarwal, R. & Morin, P. J. The claudin gene family: expression in normal and neoplastic tissues. BMC Cancer 6, 186 (2006).

55. Kominsky, S. L. et al. Clostridium perfringens enterotoxin elicits rapid and specific cytolysis of breast carcinoma cells mediated through tight junction proteins claudin 3 and 4. Am. J. Pathol. 164, 1627–1633 (2004).

56. Hough, C. D. et al. Large-scale serial analysis of gene expression reveals genes differentially expressed in ovarian cancer. Cancer Res. 60, 6281–6287 (2000).

57. Nichols, L. S., Ashfaq, R. & Iacobuzio-Donahue, C. A. Claudin 4 protein expression in primary and metastatic pancreatic cancer: support for use as a therapeutic target. Am. J. Clin. Pathol. 121, 226–230 (2004).

58. Michl, P. et al. Claudin-4: a new target for pancreatic cancer treatment using Clostridium perfringens enterotoxin. Gastroenterology 121, 678–684 (2001).

59. Curtis, C. et al. The genomic and transcriptomic architecture of 2,000 breast tumours reveals novel subgroups. Nature 486, 346–352 (2012).

60. Jacobson, A. R-Based API for Accessing the MSKCC Cancer Genomics Data Server. R package version 1.2. 5. (2015).

61. Gentleman, R. C. et al. Bioconductor: open software development for computational biology and bioinformatics. Genome Biol. 5, R80 (2004).

62. Schröder, M. S., Culhane, A. C., Quackenbush, J. & Haibe-Kains, B. survcomp: an R/Bioconductor package for performance assessment and comparison of survival models. Bioinformatics 27, 3206–3208 (2011).

63. Cochrane, W. G. The combination of estimates from different experiments. Biometrics 10, 101–129 (1954).

64. Royston, P. & Sauerbrei, W. A new measure of prognostic separation in survival data. Stat. Med. 23, 723–748 (2004).

65. Harrington, D. P. & Fleming, T. R. A Class of Rank Test Procedures for Censored Survival Data. Biometrika 69, 553 (1982).

66. Sandve, G. K., Nekrutenko, A., Taylor, J. & Hovig, E. Ten simple rules for reproducible computational research. PLoS Comput. Biol. 9, e1003285 (2013).

67. Gentleman, R. Reproducible research: a bioinformatics case study. Stat. Appl. Genet. Mol. Biol. 4, Article2 (2005).

68. Stroup, D. F. et al. Meta-analysis of Observational Studies in Epidemiology: A Proposal for Reporting. JAMA 283, 2008–2012 (2000).

